# Aging-related mechanical degradation of cortical bone is driven by microstrucural changes in addition to porosity

**DOI:** 10.1101/2023.03.01.530672

**Authors:** André Gutiérrez Marty

## Abstract

This study aims to gain mechanistic understanding of how aging-related changes in the microstructure of cortical bone drive mechanical consequences at the macroscale. To that end, cortical bone was modeled as a bundle of elastic-plastic, parallel fibers loaded in uniaxial tension, which comprised osteons and interstitial tissue. Distinct material properties were assigned to each fiber in either the osteon or interstitial fiber “families.” Models representative of mature (20-60 yrs.) bone, and elderly (60+) bone were created. Aging-related changes were modeled along three independent dimensions: (i) increased porosity, (ii) increased ratio of osteon fibers relative to interstitial fibers, and (iii) a change in fiber material properties.

The model captured decreases in modulus, yield stress, yield strain, ultimate stress, ultimate strain, and toughness with age of 14%, 11%, 8%, 6%, 20%, and 30%, respectively. In both mature and elderly bundles, rupture of the interstitial fibers drove the initial loss of strength following the ultimate point. Plasticity and more gradual rupture of the osteons drove the remainder of the response. Both the onset and completion of interstitial fiber rupture occurred at lower strains in the elderly vs. mature case.

Changes along all three dimensions were required for the model to capture aging-related decline in the strength, ductility, and toughness of cortical bone. These findings point to the importance of studying microstructural changes beyond porosity, such as the area fraction of osteons and the microconstituent material properties of osteon and interstitial tissue, in order to further our understanding of aging-related changes in bone.

## 1 Introduction

Bone tissue is a multiscale living material that fulfills important biomechanical demands, such as to support movement and protect vital organs. The highly dense type of bone tissue that encases all bones, cortical bone, contributes to whole-bone stiffness and strength [1] and is characterized by a complex hierarchical microstructure [2]. At times, cortical bone sustains mechanical loads that strain the tissue beyond its elastic regime [3]. Evidence indicates that aging impairs the ability of cortical bone to withstand these post-yield strains, and that this impairment may contribute to the increased incidence of fracture with aging [4]. In particular, degradation of mechanical properties such as strength and toughness is evident with aging [5]. However, less is clear about how aging-related changes in the microstructure of cortical bone [6] may be driving these changes in mechanical behavior at the macroscale. Since cortical bone plays a key load-bearing role in the skeleton, studying how the inelastic mechanical behavior of this tissue depends on its microstructure over the course of aging is therefore critical to understanding how and when bones fail.

The microstructural constituents of cortical bone differ from one another in their mechanical behavior, and both these behaviors and the relative amounts of these constituents have been observed to change with aging [5–8]. Mature human cortical bone is characterized by a Haversian structure that is made up of osteons encased in an interstitial bone matrix. Osteons are structures of bone tissue composed of mineralized collagen fibers that are layered parallel to each other into stacks of sheets (lamellae). These concentric lamellae are wrapped around a central pore (Haversian canal) and are bordered by a cement line. Interstitial bone tissue is the primary bone which has not yet remodeled into an osteon, as well as the fragments of osteons that remain after portions of those osteons have been remodeled. Interstitial bone tissue is more mineralized [9], stiffer by approximately 20% [6, 10–17], and less resistant to fracture [18] than the bone tissue in intact osteons. Osteons from older bones exhibit lower toughness [7], higher compressive strength [19], and similar stiffness [6, 12] and tensile strength [8], compared to those from younger bones. However, comparisons across broad age ranges are lacking due to experimental challenges pertaining to the difficulty of isolating single osteons for testing. These challenges may also contribute to the high variability in the reported data. Meanwhile, evidence suggests aging-related increases in the strength, yield stress, and stiffness, along with a decrease in toughness, of interstitial tissue [20]. The volume fraction of osteons has been shown to increase with aging [5, 21], while that of interstitial bone tissue decreases [21].

How aging-related differences in the microscale constituents of cortical bone influence the failure behavior at the macroscale is not well understood [6]. Porosity increases with aging and is strongly correlated with loss of stiffness, strength, and toughness [5, 6, 12, 21–25], but can only account for about 75% of the variation in strength and 60% in that of toughness [5, 6, 26, 27]. Evidence of aging-related changes in mineral content, which correlates with stiffness, is mixed, with some reports of increasing ash content [27] and others suggesting no change in ash content[5, 28], calcium content [5, 28, 29], or mineral-to-matrix ratio [6]. Furthermore, stiffness and strength have been shown to correlate with morphological changes in the microstructure [28], such as increasing number of osteons [5, 21, 30], decreasing area fraction of osteons [25], and increasing Haversian canal area [25, 30]. Taken together, these studies lend credence to the idea that multiple aging-related changes in morphology of the microstructure, rather than just porosity, may be important determinants of bone mechanical properties. Studies have also shown that changes in the interstitial fracture toughness [31] and mechanical integrity of the collagen network, rather than porosity [28, 32, 33], may play an important role in the toughness of cortical bone. Mechanistic interpretation of how these various morphological, compositional, and mechanical factors at the microscale, alone and in combination, affect the mechanical behavior at the macroscale is still lacking.

This study aims to understand how aging-related changes in the microstructure of cortical bone may drive mechanical consequences at the macroscale. Given the difficulty in using an experimental approach to isolate individual, aging-related changes, we used a mechanical model. With this model, we explored the effects of aging-related changes in material properties, the relative volume fraction of osteons and interstitial bone tissue, and porosity on the mechanical behavior of cortical bone.

## 2 Methods

We consider the mechanical behavior of cortical bone loaded in the longitudinal direction. We model cortical bone as a heterogeneous bundle of straight, longitudinally oriented, elasto-plastic fibers. Damage is modeled by the progressive rupture of individual fibers in the bundle. This type of one-dimensional (1D) damage model [34] of cortical bone has been used previously to analyze the nonlinearity in the tensile response of cortical bone [35, 36]. We extend that model here to incorporate plasticity as well as two fiber families, one representing osteons and the other interstitial tissue. Each fiber family has its own distributions of modulus, yield stress, and rupture stress, as depicted in **Figure 1**. Thus, the model represents heterogeneity on two scales. At the scale of the fiber bundle, the difference in mechanical behavior between the two types of microstructural constituents (osteons, interstitial tissue) is considered by varying the material properties and volume fraction of each fiber family. At the scale of individual fibers, the mechanical heterogeneity within each type of microconstituent is considered by sampling from a distribution for each fiber family.

**Figure 1:**
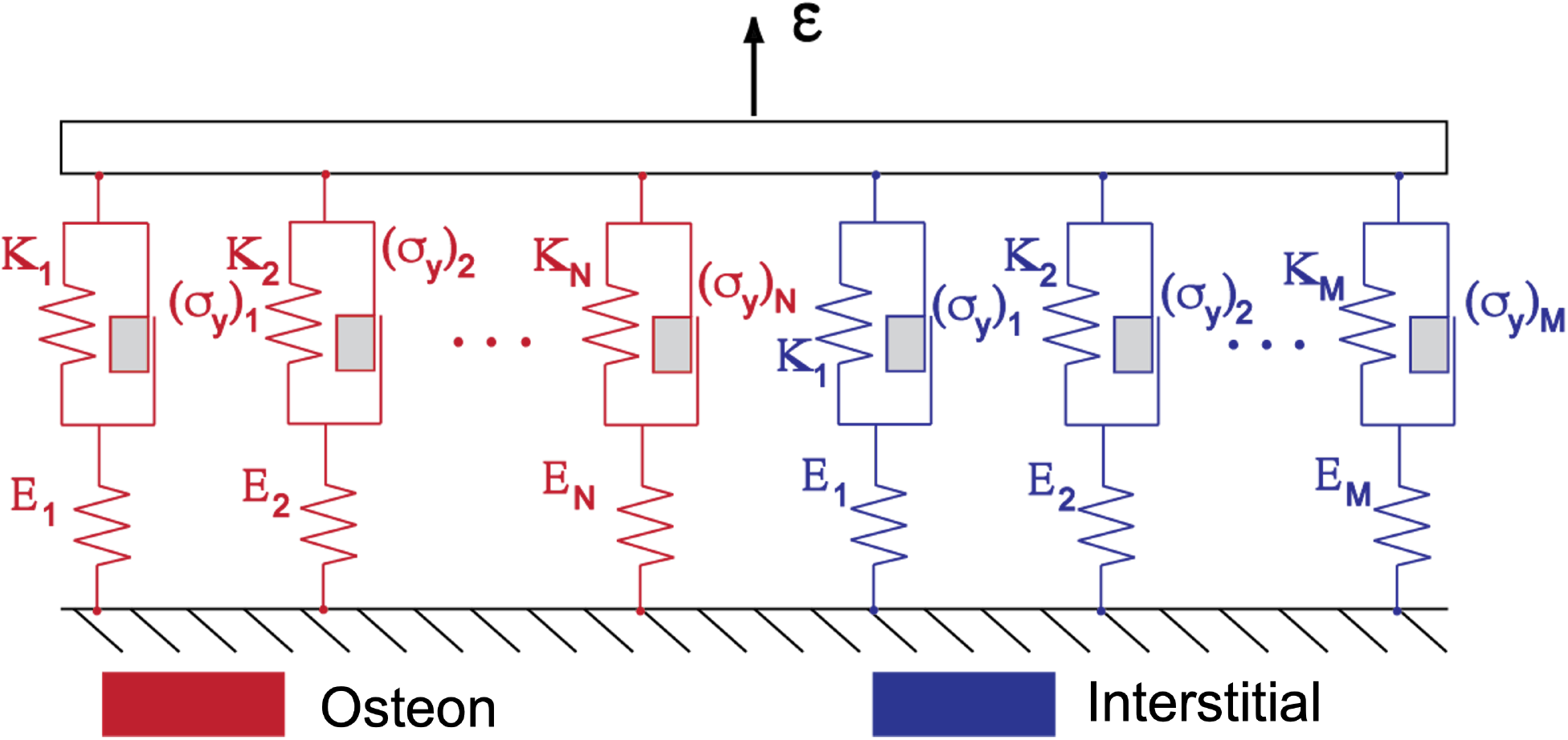
Cortical bone is modeled in a one-dimensional setup consisting of a bundle of parallel fibers loaded in strain-control. Each fiber is represented as a primary elastic spring (*E*) connected in series with a plastic element comprising a secondary spring (*K*) mounted in parallel with a Coulomb block that undergoes permanent deformation when its yield stress (*σ_y_*) is exceeded. Distinct material properties, sampled from normal distributions, are assigned to each fiber within each osteon (red) and interstitial (blue) fiber “family”.

Microstructural changes apparent with aging are represented in two different ways. First, the volume fraction of each fiber family is assigned to represent the aging-related changes in the makeup of, and porosity in, cortical bone. Second, the property distributions of the fiber families are selected to also represent aging-related changes in the material properties of each microconstituent. We assume the bone to be disease-free, and therefore assign material properties that correspond to healthy cortical bone. In this way, observed aging-related changes in the intrinsic material properties and morphology of bone tissue can be modeled through the properties of the fiber families, and the outputs of the model, the bundle’s mechanical properties, represent the macroscale mechanical behavior of cortical bone.

### 2.1 Fiber bundle model: constitutive behavior

#### 2.1.1 Bundle stress and strain

The stress state of the entire fiber bundle (*σ*) can be determined in terms of the stress 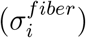 in each individual fiber *i* within the bundle. Assuming all fibers have the same crosssectional area (*A^fiber^* = constant), the bundle stress (*σ*) is given by the total force (*F_TOT_*) and cross-sectional area (*A_TOT_*):

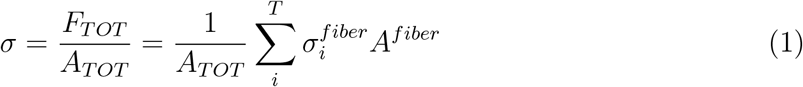

Since all fibers have constant cross-sectional area, the volume fraction of each fiber family (*ϕ^OST^*, *ϕ^INT^*) can be determined by the number of fibers belonging to that family:

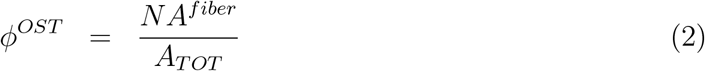

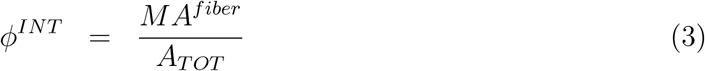

where *N* refers to the total number of fibers in the osteon (OST) fiber family and *M* the total number of fibers in the interstitial (INT) family. The porosity, *ϕ^PORE^*, is defined as *ϕ^PORE^* = 1 – *ϕ^OST^* – *ϕ^INT^*. Hence, the bundle stress is now given by:

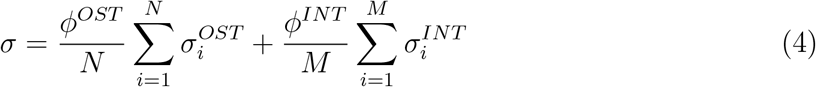

We assume the fibers to be loaded in parallel such that all fibers in the bundle are strained uniformly. Therefore, the strain on each fiber 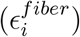 is equal to the bundle strain (*ϵ*). With this assumption in mind, the simulated loading in the model is strain-controlled.

#### 2.1.2 Mechanical behavior of an individual elastic-plastic fiber

Each fiber is modeled as elastic-plastic with linear hardening prior to rupture. The fiber stress is fully characterized by the fiber strain, accumulated plastic strain in the fiber, and the following material properties, where the subscript *i* is omitted for clarity:

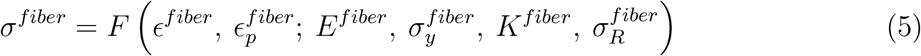

We discretize the load history over time into load steps – assuming equilibrium at each load step – 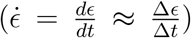, and assuming that the relevant state variables from the prior load step are known, such that *σ_n_* = *σ^fiber^*, *ϵ_n_* = *ϵ*, and 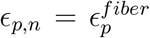. Variables with the subscript *n* refer to the previous load step (known), while those with subscript of *n* + 1 refer to values at the current load step (unknown). We drop the superscript “fiber” for the variables below, since these all correspond to a single fiber. To determine the stress increment in a given fiber due to a prescribed strain step Δ*ϵ*, we use the trial stress 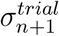, which is the predicted stress for a given strain corresponding to the elastic modulus of the fiber; i.e., the stress corresponding to an elastic step [37]:

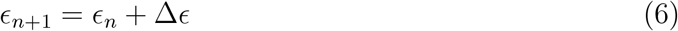

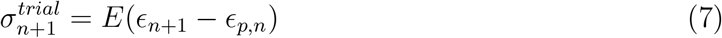

A yield function 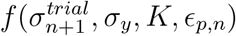 is used to determine if the fiber has yielded. If so, then plastic strain, *ϵ_p_*, will be updated accordingly. Linear strain-hardening and tensile loading are assumed, with |*ϵ_p_*| as the hardening parameter[37]. The yield function is therefore:

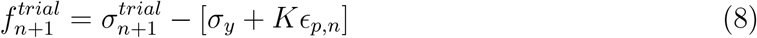

The deformation and stress in the fiber at the given strain increment can then be determined by the sign of the resulting value for the yield function, where:

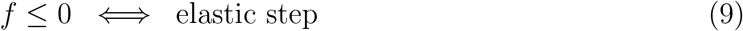

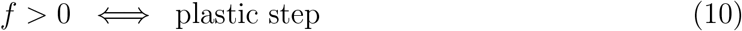

##### Elastic step 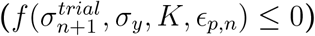

In this case, the fiber stress is computed directly based on the elastic modulus of the material, and only the elastic strain is considered:

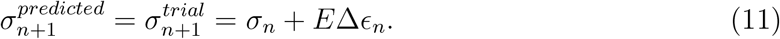

##### Plastic step 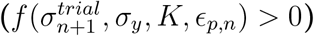

In this case, the fiber yields, and the stress is updated according to a return mapping algorithm [37] in which the trial stress is projected onto, or mapped back (“returned”), to the yield surface. The plastic strain is updated in terms of the plastic slip (Δ*γ*):

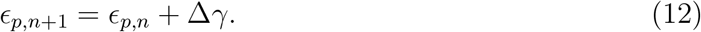

The plastic slip is the increase in plastic strain due to the change in the yield surface due to hardening:

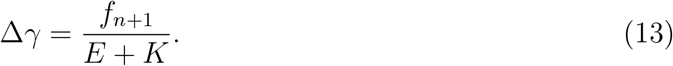

Hence, the fiber stress can be predicted by

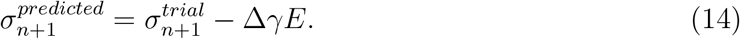

##### Check for rupture

Regardless of yielding, the fiber will rupture if its predicted stress exceeds its rupture strength (*σ_R_*). First, the predicted fiber stress 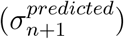 is computed from Eq. 11 or 14, and the corresponding value is used in Eq. 15 below:

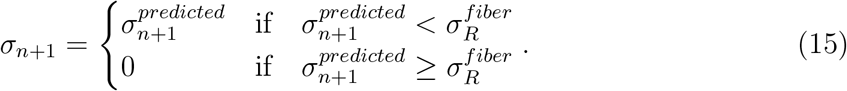

If 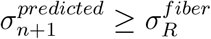, the fiber has ruptured and so its modulus is diminished accordingly,

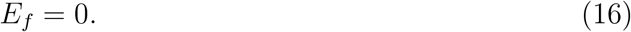

#### 2.1.3 Assignment of material properties

The moduli and rupture strengths of each fiber family were assumed to follow normal distributions. The parameters characterizing each property’s distribution are listed in Table 1. The choices of their values are discussed below. The manner by which properties are assigned and distributed has significant ramifications on the bundle behavior. Some of the consequences surrounding the types of distributions and their types are presented in the appendix. The properties of each individual fiber were randomly sampled from the assumed distributions assigned to its fiber family (i.e., osteon, interstitial).

**Table 1:**
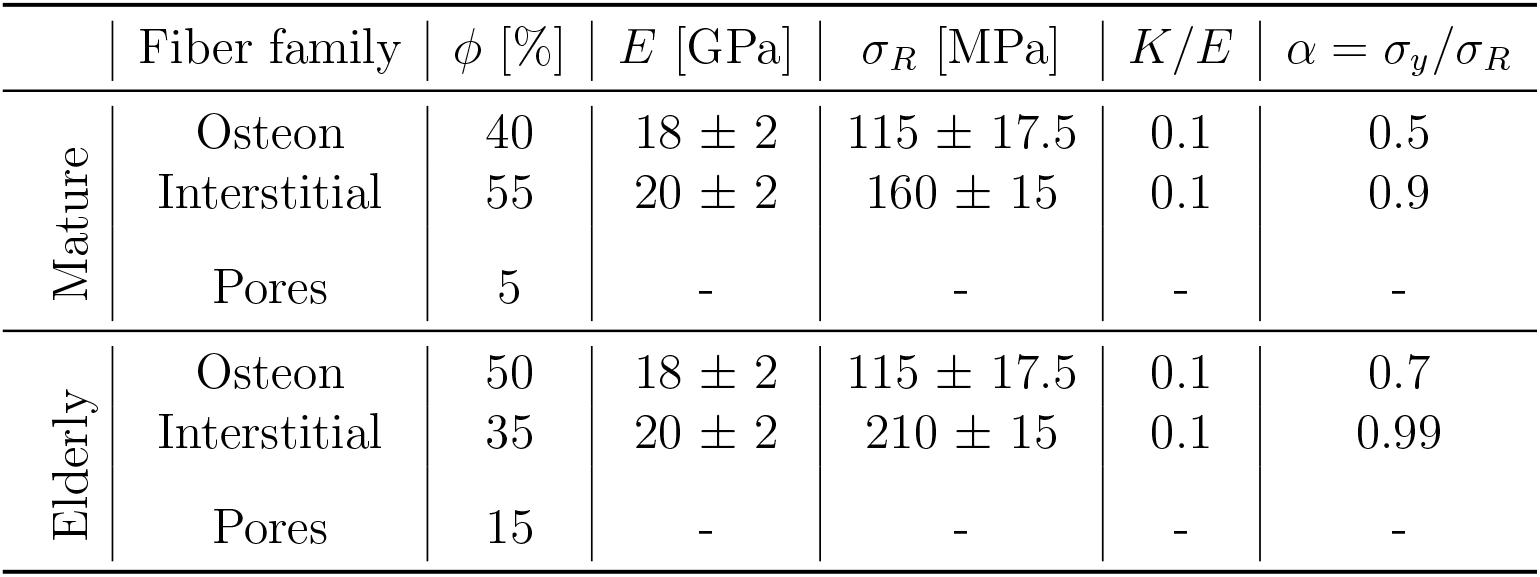
Model inputs for mature bone (baseline case) and elderly bone. Mean ± stdev are specified for prescribed material property distributions.

Data for osteon Young’s modulus were available from both nanoindentation studies and tensile tests of individual osteons. Nanoindentation studies, in which a nanoindenter was used to measure the elastic moduli within individual lamellae, reported values of 14-22 GPa [10, 12, 14, 16, 38, 39] for the tissue modulus of hydrated, fully mineralized osteons sampled from healthy human cortical bone. These values are in line with moduli reported from tensile tests on whole osteons (6-22 GPa) [8]. Since interspecimen (osteon-to-osteon) coefficient of variation was estimated to be approximately 10% on average based on nanoindentation tests [39], and 50% at most based on tensile tests [8], we base our modulus values primarily on nanoindentation data. Therefore, we assume osteon moduli fall into a normal distribution centered at 18 GPa with coefficient of variation of ~ 10%.

Data for osteon inelastic behavior were available from tensile tests of individual osteons. Taking into account previously measured average dimensions of whole osteons and their corresponding Haversian canals [25], the ultimate strengths reported from tensile tests on osteons were adjusted to estimate the effective ultimate stress of the bone tissue within the osteon alone (i.e., excluding the Haversian canal area). Those tensile experiments reported osteon ultimate stress in the range of 88-135 MPa, and also reported that the proportional limit (i.e., the transition to nonlinear mechanical behavior as measured on a representative stress-strain curve) was approximately half of its ultimate stress. Accounting for the fraction of osteons occupied by Haversian canals gives a tissue ultimate stress estimate in the range 93-144 MPa. Therefore, we assumed osteon rupture strengths fall into a normal distribution centered at 115 MPa with coefficient of variation of 15%, and set *α^OST^* = *σ_y_/σ_R_* = 0.5. The plastic modulus (*β^OST^* = *K/E* = 0.1) was obtained by computing the secant modulus of the plastic region of the stress-strain curve shown by Ascenzi et al. [8] of a fully mineralized, wet osteon. These four properties, *α^OST^*, *β^OST^*, *E*, *σ_R_*, fully determine the bilinear stress-strain curve (Figure 2) modeling a given osteon fiber.

**Figure 2:**
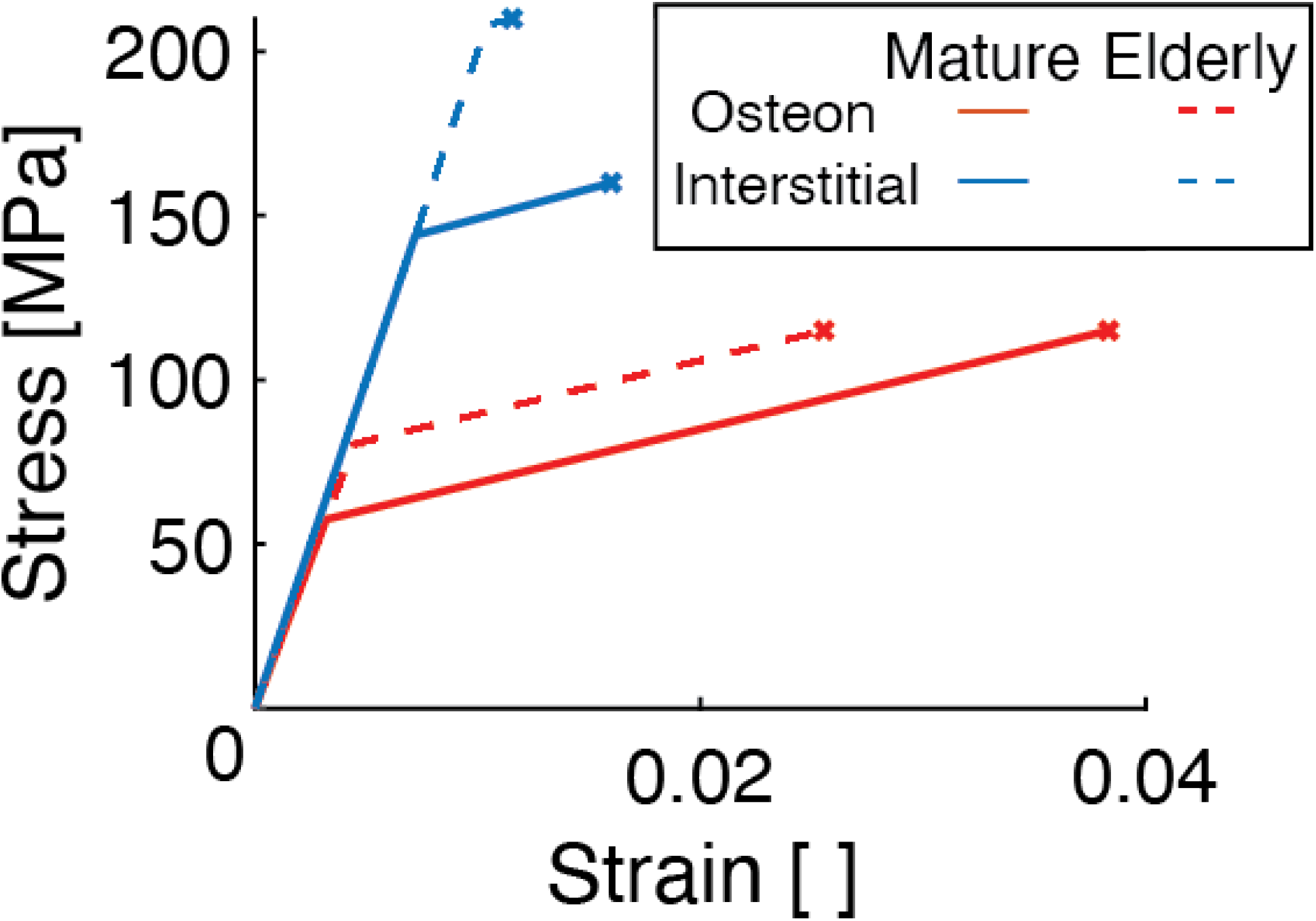
The stress in a representative mature osteon fiber (solid red), a mature interstitial fiber (solid blue), an elderly osteon fiber (dashed red), and an elderly interstitial fiber (dashed blue). “X” marks the failure point for each fiber.

Data for interstitial Young’s modulus were available from nanoindentation studies and tensile tests of primary bone tissue. Nanoindentation studies report moduli in the range of 15-26 GPa [6, 12, 14, 16, 39] for samples of interstitial bone tissue. These values were in line with values reported from tensile tests on primary bone tissue (about 20 GPa) [40]. Inter-specimen variability of interstitial tissue was found to be under 10% in those studies. Therefore, we assume interstitial moduli fall into a normal distribution centered at 20 GPa with a coefficient of variation of 10%.

The inelastic behavior of interstitial tissue was specified based on the tensile data on primary bone tissue. Since no data directly analogous to the tensile tests of individual osteons exists for interstitial bone tissue, we assumed that measurements of primary bone mechanical properties [40] are representative of interstitial bone. Based on these data, we assumed interstitial rupture strengths fall into a normal distribution centered at 160 MPa with standard deviation of 15 MPa. Taking into account the highly mineralized nature of interstitial bone tissue as compared to osteon tissue [41], which results in a lower resistance to fracture than that for osteons [7, 18, 23], the yield stress was set to be only slightly lower (10%) than the corresponding strength of each fiber (i.e., *α^INT^* = *σ_y_/σ_R_* = 0.9). This assumption allows only minimal post-yield deformation. The plastic modulus was estimated by assuming the toughness to be 20% lower than that of osteon tissue, resulting in *β^INT^* = *K/E* = 0.1; see Figure 2.

Aging-related changes in each of the two microconstituent tissues are observed to be different, and therefore modeled accordingly. The previously mentioned values for osteon and interstitial elastic and inelastic parameters, summarized in Table 1, represent the baseline case of healthy, mature (20-60 years of age) cortical bone. Different values are chosen to reflect aging-related microstructural and material changes to establish the model of elderly cortical bone (60+ years of age), and are done so independently of a change in porosity.

Aging of osteon fibers was modeled primarily by reducing their toughness in addition to increasing their volume fraction with respect to interstitial fibers. As mentioned previously, values of osteon elastic modulus [8, 12] and osteon strength [8] measured from a population of elderly samples were not distinct from those taken from mature samples. Therefore, the property distributions corresponding to moduli and rupture strengths of osteon fibers were set to be the same in the mature and elderly bone models. Two options exist for reducing fiber toughness to model reported measurements [7]: increasing *β* or increasing *α*. We chose the latter and raised *α^OST^* to 0.7 (Figure 2) assuming a reduction in toughness of 25% and known *α^OST^, E, σ_R_*.

Aging of interstitial fibers, on the other hand, was modeled by reducing their toughness and increasing their rupture strength, in addition to the aforementioned change in volume fraction. Preliminary data reported in [20] suggest that the ultimate strength and toughness of interstitial tissue from older subjects are approximately 30% higher and 30% lower, respectively, than that from younger subjects. Therefore, the mean strength of interstitial fibers was increased to 210 MPa [40], and *α^INT^* was raised to 0.99.

Using this fiber bundle model, we characterized elderly cortical bone in terms of changes in model inputs along the following three dimensions: morphological aging, porosity increase, and material aging (Table 2). Morphological aging refers to the increase of the volume fraction of osteon fibers (*ϕ^OST^*) with respect to that of interstitial fibers (*ϕ^INT^*). Porosity increase refers to the reduction of the total solid volume fraction (comprising osteon and interstitial fibers), thus increasing the porosity (*ϕ^PORE^*) in the model. Finally, material aging refers to the aforementioned changes in osteon (*α^OST^*) and interstitial (*α^INT^*,(*σ_R_*)*^INT^*) inelastic parameters.

**Table 2:** Model inputs changed between mature (baseline), elderly, and additional cases; 0 signifies no change, while bold **1** signifies an aging-related change.

| Change label | Morphological | Porosity | Material |
| --- | --- | --- | --- |
| Properties changed ($\Delta$) | $(\phi^{OST} : \phi^{INT})$ | $(\phi^{PORE})$ | $\begin{pmatrix} (\sigma_R)^{INT} \\ \alpha^{OST} \\ \alpha^{INT} \end{pmatrix}$ |
| (1) Baseline (mature bone) | 0 | 0 | 0 |
| (2) Elderly bone | **1** | **1** | **1** |
| (3) Morphological aging | **1** | 0 | 0 |
| (4) Porosity increase | 0 | **1** | 0 |
| (5) Material aging | 0 | 0 | **1** |
| (6) Morphological aging & porosity increase | **1** | **1** | 0 |
| (7) Morphological & material aging | **1** | 0 | **1** |
| (8) Material aging & porosity increase | 0 | **1** | **1** |

#### 2.1.4 Brittle model and the role of plasticity

Simulations corresponding to the volume fraction (*ϕ*), modulus (*E*), and strength (*σ_R_*) parameters listed in Table 1 were repeated with a brittle fiber model, in which all fibers were considered perfectly elastic, for both mature and elderly cortical bone. The comparison of the predictions of the perfectly brittle model to those of the elastic-plastic model will reveal the effect of the plastic behavior of microstructural constituents (i.e. osteon and interstitial bone tissue) on the overall behavior of the model.

In this brittle model, the mechanical behavior for each fiber was fully defined by its modulus and rupture strength; yield stress (*σ_y_*) was equal to the rupture stress (*α* = 1) and plastic modulus (*K*) was undefined. As established for the elastic-plastic model discussed above, material properties for each fiber were randomly sampled from a Gaussian distribution according to the parameters presented in Table 1.

#### 2.1.5 Model Outputs

For each bundle modeled, *σ* was computed at each strain step, and the stress-strain (*σ*-*ϵ*) curve was used to quantify key aspects of the overall mechanical response. The yield point (*ϵ_y_,σ_y_*) was identified using a 0.2% offset in strain, ultimate stress (*σ_ult_*) as the highest stress sustained by the bundle, ultimate strain (*ϵ_ult_*) as the strain at the ultimate stress, and toughness (*U_T_*) as the area under the stress-strain curve.

### 2.2 Model cases

The multi-scale nature of the fiber bundle model enables a direct examination of the effects of changes to individual inputs (material properties, volume fraction, etc.) on the macroscopic mechanical behavior to understand the respective contributions of these changes. In total, eight cases were considered: the baseline (Case 1: mature bone) and elderly (Case 2) models already described, and six cases that isolate one or two changes at a time (Table 2). First, only the ratio of volume fractions of osteon fibers to interstitial fibers was increased proportionate to the change in areal osteon density observed with aging [21, 24] (Case 3: morphological aging). Secondly, only porosity was increased (Case 4: porosity increase). Then, changes in material properties were isolated, by only increasing the interstitial average rupture strength, interstitial yield stress, and osteon yield stress (Case 5: material aging). Finally, cases 6-8 capture pairs of these independent changes.

### 2.3 Sensitivity analysis

The intra-specimen variability of, for example, Young’s modulus is captured in our model by drawing Young’s modulus values from from a distribution with specified mean and standard deviation. Here we capture inter-specimen variability by considering the sensitivity of our model predictions to variability, or uncertainty in, for example, the mean value of Young’s modulus specified in the model. Therefore, a local sensitivity analysis was conducted to investigate the effects of biological variation of, or our uncertainty in, model inputs on the bundle’s macroscale mechanical properties. The sensitivities of ultimate stress, yield stress, ultimate strain, yield strain, and toughness to the moduli (mean *E^OST^, E^INT^*), strengths (mean (*σ_R_*)^*OST*^, (*σ_R_*)^*INT*^), yield stress ratios (*α^OST^*, *α^INT^*), and plastic modulus ratios (*β^OST^*, *β^INT^*), as well as to osteon and pore volume fractions (*ϕ^OST^*, *ϕ^PORE^*), were evaluated. For each of these macroscale mechanical properties, the change in the property’s value relative to a change in each one of the model inputs was computed, then normalized by the ratio of standard deviations [42]:

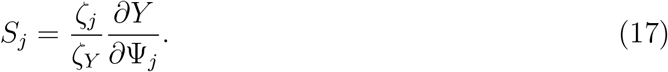

Here, *Y* represents each mechanical property (i.e.: *E, ϵ_y_, σ_y_, ϵ_ult_, σ_ult_, U_T_*), Ψ_*j*_ represents the *j^th^* input parameter (e.g., osteon modulus, interstitial modulus, osteon strength,…), *ζ_j_* is the expected standard deviation in the input Ψ_*j*_, and *ζ_Y_* is the variation in the outputs, expressed as a standard deviation and given by 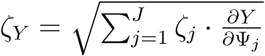, with *J* = 10 equalling the total number of model inputs. We note that 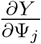 was computed by finite differences at the nominal value, Ψ_*j*_. We also note that the standard deviations initially used to define material property distributions (*E, σ_R_*) are representative of the expected variation across a specimen population and are therefore considered to be representative of the natural biological variation between specimens within the sampled population in this sensitivity analysis. We consider the range of possible *β* values as log-normal, and characterize our uncertainty in β as the standard deviation of log_10_ *β*.

## 3 Results

The stress-strain curves for the fiber bundle models representing mature and elderly cortical bone both exhibited an elastic region followed by apparent hardening and then a progressive loss of strength (Figure 3). Loss in macroscale toughness and strength of the elderly model compared to mature was evident. Direct numerical comparison showed decreases with age in modulus, yield stress, yield strain, ultimate stress, ultimate strain, and toughness by 14%, 11%, 8%, 6%, 20%, 30%, respectively (Table 3). The amount of stress borne by each fiber family over the course of loading also changed with age (Figure 3). Despite the increased strength and equivalent stiffness of the interstitial fibers in the elderly case, the decreased volume fraction of these fibers resulted in a decrease in the proportion of stress borne by them (dashed blue) and thus, decreased ultimate strength at the macroscale. Concomitant with a decrease in volume fraction of interstitial fibers is an increase in volume fraction of osteon fibers, which is responsible for osteon fibers carrying an increased portion of the total stress in the elderly bone model. Figure 3 also shows a decrease in toughness for the elderly cortical bone model (black dashed) as compared to the mature model (black solid). This overall reduction in toughness is due in part to a combination of the decrease in osteon toughness and its larger volume fraction.

**Figure 3:**
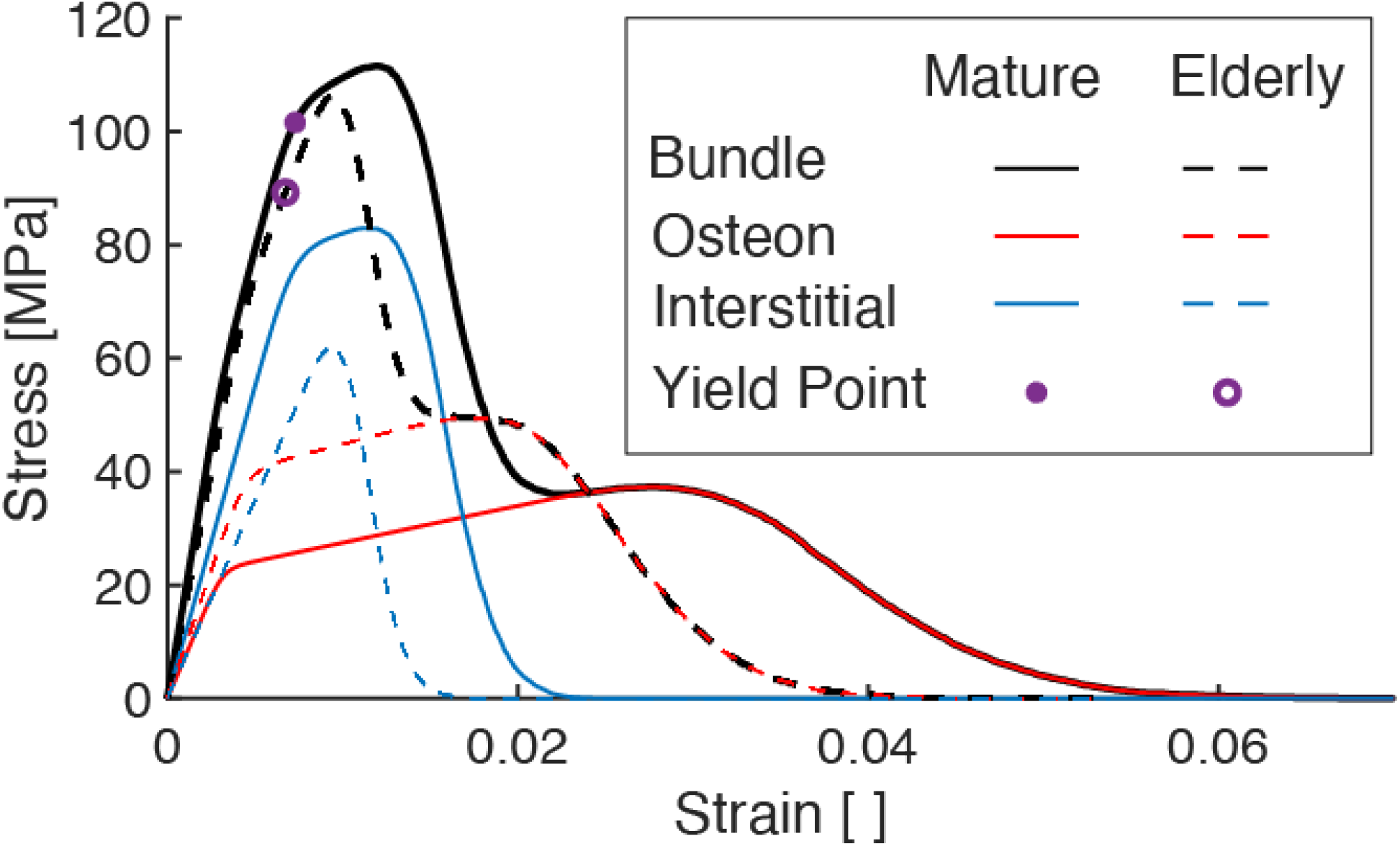
Stress-strain curves of elastic-plastic fiber bundle model comparing the mechanical behavior of the mature bone and elderly bone model cases. The blue (and red, respectively) curves show the partial stress borne by the interstitial (and osteon, respectively) fibers. The sum of the partial stresses give the total stress (black curves).

**Table 3:** Model outputs for bundle mechanical properties corresponding to various cases

| Case | $E$<br>[MPa] | $\epsilon_y$<br>$\cdot 10^{-2}$ | $\sigma_y$<br>[MPa] | $\epsilon_{ult}$<br>$\cdot 10^{-3}$ | $\sigma_{ult}$<br>[GPa] | $U_T$<br>[MJ/m<sup>3</sup>] |
| --- | --- | --- | --- | --- | --- | --- |
| (1) Baseline (mature bone) | 18.2 | 7.29 | 101 | 1.18 | 113 | 2.34 |
| (2) Elderly bone | 16.0 | 6.74 | 89.3 | 0.95 | 106 | 1.63 |
| (3) Morphological aging | 17.9 | 6.57 | 86.2 | 1.21 | 99.7 | 2.54 |
| (4) Porosity increase | 16.3 | 6.46 | 85.5 | 1.18 | 100 | 2.08 |
| (5) Material aging | 18.2 | 8.79 | 130 | 0.94 | 133 | 1.67 |
| (6) Morphological aging<br>& porosity increase | 16.0 | 5.78 | 71.2 | 1.21 | 88.6 | 2.28 |
| (7) Morphological<br>& material aging | 17.9 | 7.83 | 110 | 0.98 | 124 | 1.81 |
| (8) Material aging<br>& porosity increase | 16.3 | 7.43 | 104 | 0.94 | 120 | 1.48 |
| Brittle mature | 18.2 | 6.5 | 86.6 | 0.57 | 91.6 | 0.51 |
| Brittle elderly | 16.0 | 5.90 | 74.4 | 0.54 | 76.0 | 0.58 |

As an individual fiber is loaded, it may be in any one of 3 states: elastic, yielded, or ruptured. Quantifying the fraction of fibers in each of these states for each of the osteon and interstitial fibers revealed the extent of yielding and/or rupture in each family as the mature and elderly fiber bundles were strained (Figure 4). The three vertical lines in Figure 4 located at the yield strain, ultimate strain, and the strain at which the last interstitial fiber ruptured, provide visual guides as to the fractions of fibers in each state at these three key points in loading. From these results, it is evident that the apparent hardening of the mature bundle is due to plasticity in both fiber families, whereas that in the elderly bundle is predominantly due to plasticity in the osteons only. In both mature and elderly bundles, rupture of the interstitial fibers drives the initial loss of strength following the ultimate point, and the plasticity and more gradual rupture of the osteons, which deform over a longer strain range than interstitial fibers, drive the remainder of the stress-strain response.

**Figure 4:**
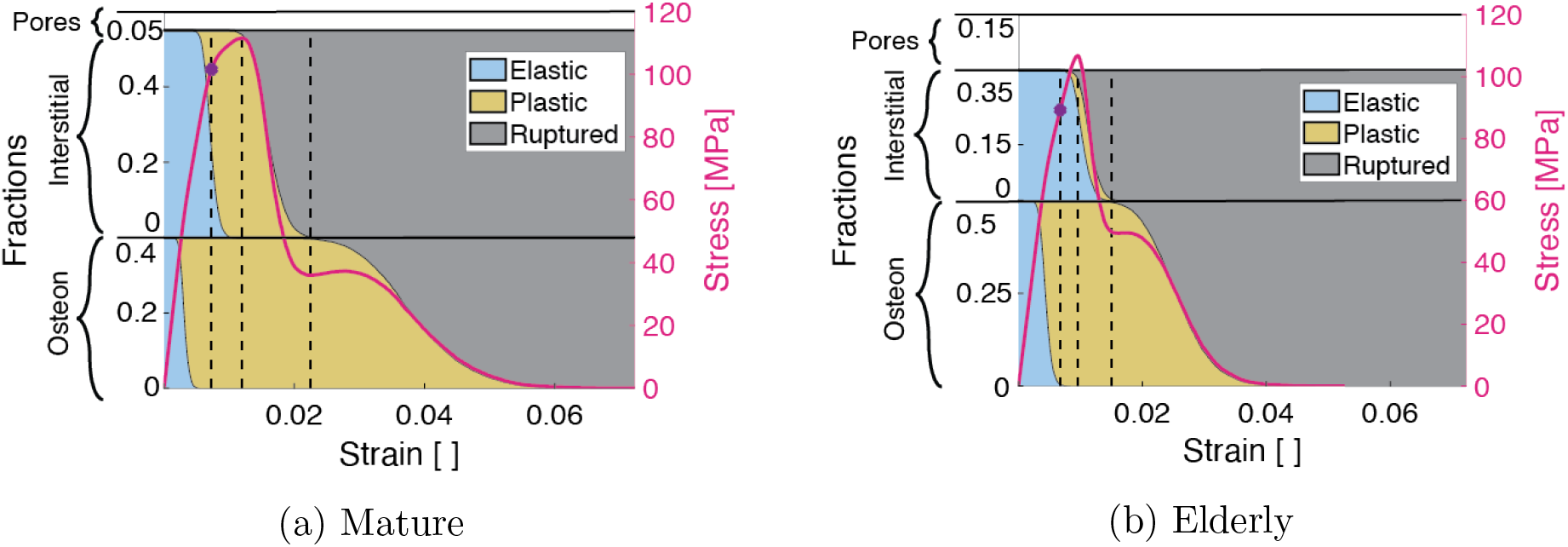
Fractions of osteon (bottom) and interstitial (top) fibers separated into elastic (blue), yielded (yellow), and ruptured (gray) regimes at each bundle deformation step. The stress-strain curve corresponding to each case (mature on the left and elderly on the right) is overlayed on the area fractions. Dotted black lines indicate the fraction of fibers pertaining to key regimes of the bundle mechanical behavior at the yield strain, ultimate strain, and the strain at which all interstitial fibers have ruptured.

However, both the onset and completion of interstitial fiber rupture occur at lower strains in the elderly vs. mature case. The reduction in strain over which interstitial fibers yield prior to rupture contributes to aging-related decreases in ultimate strain and toughness.

Additional model cases that explored changes in individual inputs (Figure 5) indicate that the differences in mechanical behavior between mature (Case 1) and elderly (Case 2) cases are the result of changes in all three aging dimensions considered: morphological aging (increase in *ϕ^OST^/ϕ^INT^*), material aging, and porosity increase. Either morphological aging alone (Case 3) or an increase in porosity alone (Case 4) captured nearly all of the decline in yield stress, yield strain, and ultimate stress in elderly compared to mature bone. However, these two cases differ in that morphological aging alone (Case 3) caused toughness to increase, which demonstrates that an increasing density of osteons has an effect opposite to increasing porosity. Material aging by itself (Case 5), or combined with either morphological aging (Case 7) or porosity decrease (Case 8), captured the decline in ultimate strain and toughness, but led to increases of the strength parameters. Combing morphological aging and porosity increase (Case 6), on the other hand, predicted a decrease in the strength parameters, but left ultimate strain and toughness largely unaffected. These results suggest that simultaneous changes in all three dimensions of aging are required to capture the full macroscale behavior of elderly bone.

**Figure 5:**
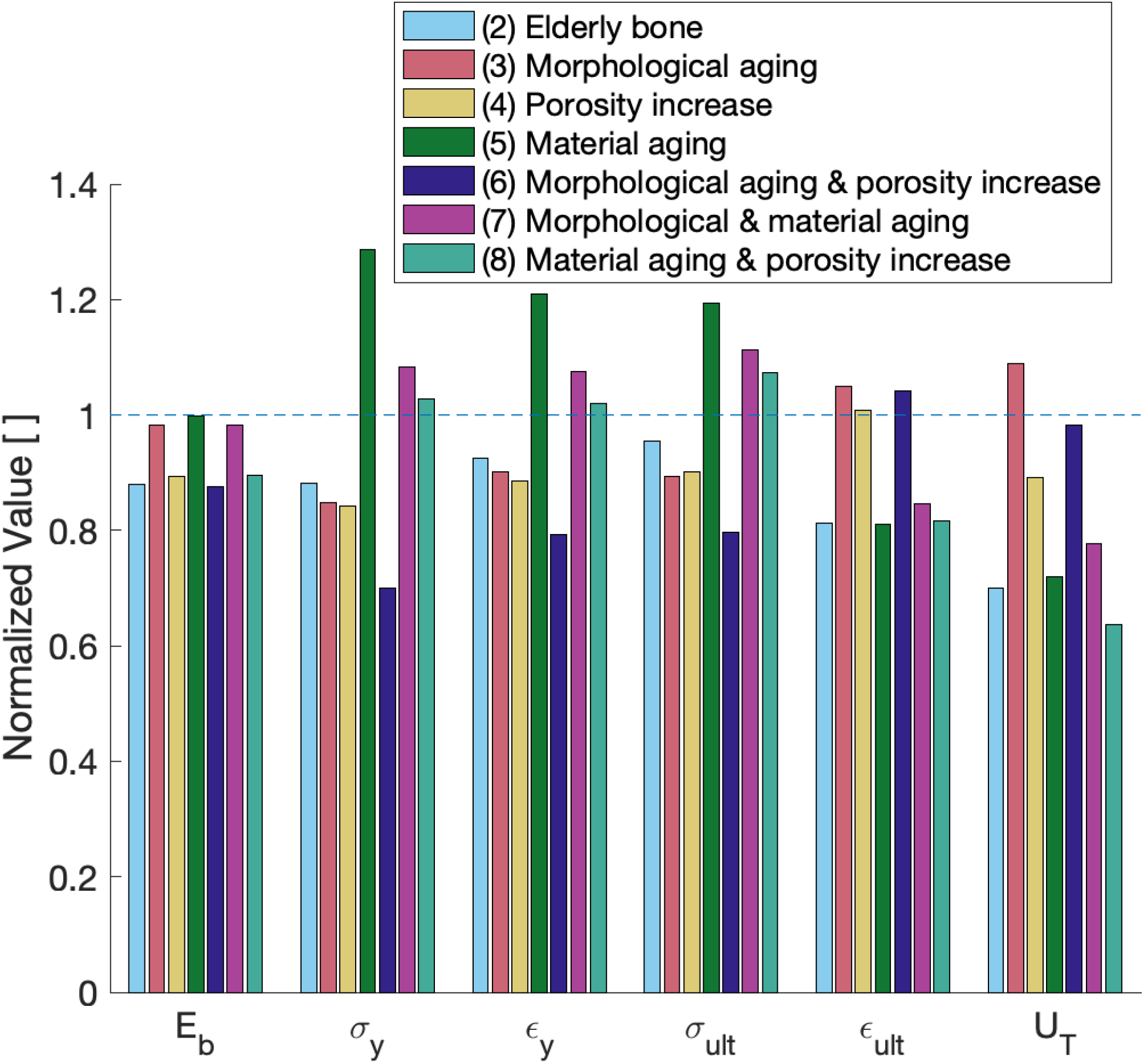
Model output values for various model cases (as listed in Table 2) normalized to the baseline case (mature bone). Aging-related include changes in material properties, increased porosity, and morphological (increased volume fraction of osteons with respect to interstitial fraction).

The brittle models, which lacked plasticity at the microstructural scale, demonstrate some but not all of these changes in mechanical properties with aging (Figure 6). While the fibers in this model were all brittle, the bundle exhibits nonlinear behavior due to a progressive accumulation of damage arising from the heterogeneity in material properties of all the fibers. The brittle model predicted decreases with age in modulus, yield stress, yield strain, ultimate stress, and ultimate strain of 3%, 36%, 3%, 34%, and 13%, respectively, along with an increase of 15% in toughness (Table 3). Compared to the plastic model results (Table 3, Figure 3), the brittle model showed larger decreases in yield and ultimate stress, but an increase in toughness.

**Figure 6:**
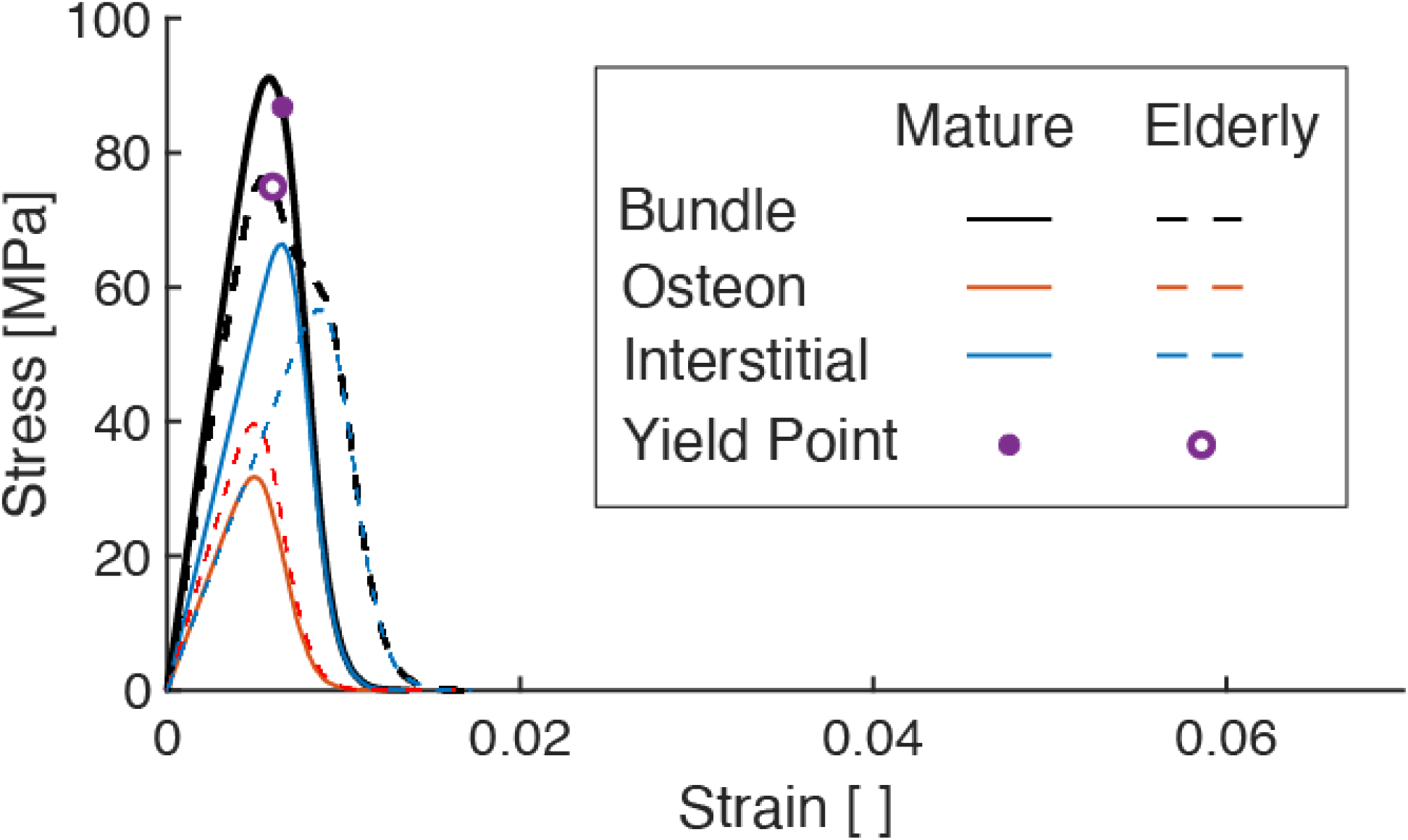
Stress-strain curves of elastic fiber model comparing the mechanical behavior of mature (M) and elderly (E) bone model cases

Among the input parameters considered, the elastic-plastic model results were most sensitive to changes in the yield and post-yield properties of both the osteon and interstitial fiber families (Figure 7). Chief among these properties were the plastic moduli of both fiber families and, to a lesser extent, the yield stresses. The plastic modulus and yield stress of the interstitial fibers (controlled by *β^INT^* and *α^INT^*, respectively), and the osteon fibers (controlled by *β^OST^* and *α^OST^*, respectively) were the most influential inputs to the post-yield behavior of the model. For example, the sensitivities of the ultimate stress and toughness to *β^OST^* and *β^INT^* were more than 2.2-fold greater than those to all other inputs, and the same was true for the sensitivity of ultimate strain to *α^INT^* and β^INT^. Similarly, the sensitivities of yield stress and yield strain to *α^OST^*, *α^INT^*, and *β^OST^* were about 2.4-fold greater than those for all other inputs. In contrast, model outputs exhibited low sensitivities across the board to osteon modulus (*E^OST^*), osteon rupture strength 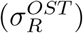, osteon volume fraction (*ϕ^OST^*), interstitial modulus (*E^INT^*), and porosity (*ϕ^PORE^*). To facilitate comparison of sensitivities among outputs, we computed the coefficients of variation, defined as the standard deviation in output, *ζ_Y_*, divided by the output value for the baseline cases (**Table 4)**. Toughness and ultimate strain had much greater coefficient of variation (296% and 250%, respectively) than did all the other outputs (30%, on average), indicating that these two outputs were the most sensitive to uncertainty in input values.

**Figure 7:**
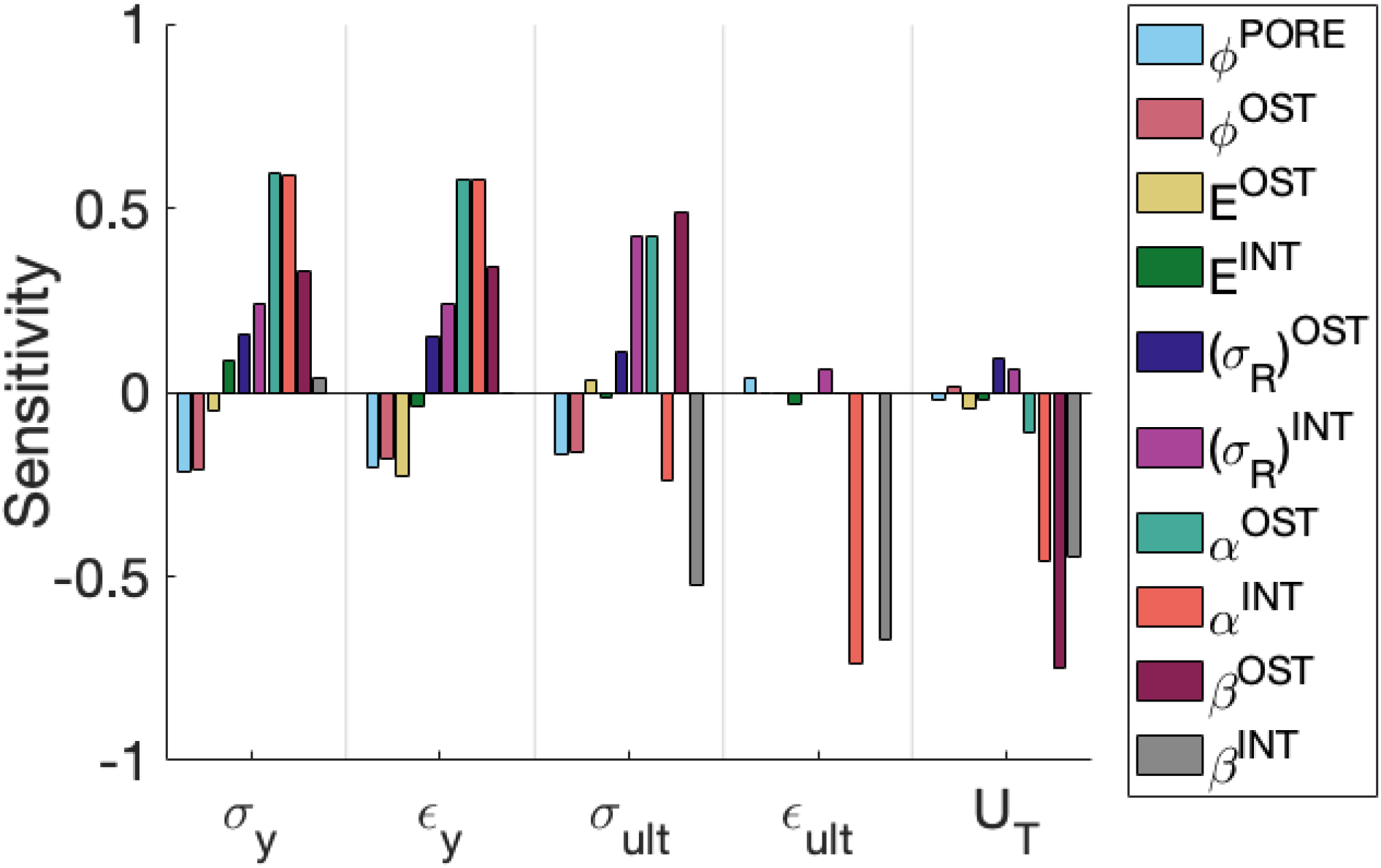
Bar chart showing sensitivity of model outputs to various model inputs (according to a finite difference evaluation of Eq 17)

**Table 4:** Estimated uncertainty or variance in input values (expressed as its standard deviation) and resulting variation in outputs.

| Inputs ($\Psi_j$) | $\zeta_j$ | Mean | CoV [%] | | Output (Y) | $\zeta_Y$ | Mean | CoV [%] |
| --- | --- | --- | --- | --- | --- | --- | --- | --- |
| $\phi^{PORE}$ | 2.5% | 5% | 50 | | $\sigma_y$ [MPa] | 29.9 | 101 | 30 |
| $\phi^{OST}$ | 5% | 40% | 12.5 | | $\epsilon_y$ | 0.0016 | 0.0073 | 22 |
| $E^{OST}$ [GPa] | 2 | 18 | 11 | | $\sigma_{ult}$ [MPa] | 41.7 | 113 | 37 |
| $E^{INT}$ [GPa] | 2 | 20 | 10 | | $\epsilon_{ult}$ | 0.0294 | 0.0118 | 250 |
| $(\sigma_R)^{OST}$ [MPa] | 15 | 115 | 13 | | $U_T$ [MPa] | 6.93 | 2.34 | 296 |
| $(\sigma_R)^{INT}$ [MPa] | 17.5 | 160 | 11 | | | | | |
| $\alpha^{OST}$ | 0.25 | 0.5 | 50 | | | | | |
| $\alpha^{INT}$ | 0.25 | 0.9 | 28 | | | | | |
| $\log_{10}(\beta^{OST})$ | 1 | -1 | 100 | | | | | |
| $\log_{10}(\beta^{INT})$ | 1 | -1 | 100 | | | | | |

## 4 Discussion

In this study, we implemented a one-dimensional mechanical model to understand how aging-related changes in the microstructure of cortical bone drive mechanical consequences at the macroscale. Simulations using the mature and elderly models with elastic-plastic fibers demonstrated decreased stiffness, yield stress and yield strain, ultimate stress and ultimate strain, and, most of all, toughness with age. Results of the cases that simulated only subsets of aging-related changes in inputs indicated that the deficits in macroscale mechanical behavior of the elderly model resulted from morphological and material changes at the microscale to a level on par with the effects of increased porosity. Morphological aging alone captured approximately all of the decline in yield and ultimate stress observed when comparing the elderly case to mature bone, and material aging alone captured the decline in ultimate strain. Interestingly, material aging alone, but not an increase in porosity, reduced toughness to a value on par with that seen for elderly bone, suggesting that a change in microstructural material properties may be a route to loss of toughness in cortical bone with age. The model with brittle fibers, however, predicted an increase in toughness, counter to experimental findings [5, 20, 43–45]. The expected decrease in toughness is found when fiber plasticity is incorporated into the model, indicating that plasticity of the osteons and interstitial tissue, rather than only damage (fiber rupture), may play a key role in the aging-related decline in macroscopic cortical bone toughness. Collectively, the findings of this study illustrate that multiple facets of aging-related changes at the microscale in cortical bone may affect the macroscale behavior, via the effects of these changes on the onset and progression of yielding and rupture of osteons and interstitial tissue.

While simpler in nature than two- or three-dimensional models, our model results confirm prior findings that changing microstructural material properties, in addition to porosity, are reflected in the macroscopic, aging-related decrease in toughness [6, 46–48] due potentially to reduced ductility of the microconstituents [48]. The heterogeneity modeled here allows the estimation of the contributions of individual cortical bone microconstituents (i.e., osteon and interstitial tissue), and the effect of aging-related changes in these microconstituents, that was not possible with earlier models [49, 50]. With these microstructural material properties in hand, our baseline model successfully predicted the mean elastic modulus of cortical bone [51], yield strain, ultimate stress, and ultimate strain [6], and toughness [52] within one standard deviation, as well as yield stress [6] within 3 standard deviations of cortical bone. Furthermore, the difference in material properties between microconstituents captured by our model implies that the aging-related decrease in ultimate strain arose from changes in the interstitial material properties (Figure 4). While more complex than other one-dimensional fiber bundle models [34, 35], incorporating plasticity into the fibers enabled us to observe the well documented, aging-related decrease in toughness of cortical bone [5, 52].

This study has some limitations. First, the model is only one-dimensional. Any aging-related changes in the 2D or 3D geometry (e.g., increased circularity of osteons [53–55]) of the microconstituents, are not reflected in this model. The model does not include any changes in cross-sectional area over the length of the fibers, or any change in the mechanical behavior resulting from a change in the three-dimensional morphology of the bone tissue (e.g., misalignment of the osteon with the longitudinal or transverse axis and/or with the accompanying Haversian canal [19]). Since quasi-static loading is assumed, rate dependence of cortical bone, and how that might change with aging, is not considered. Porosity in the model represents only that contributed by Haversian canals in osteons; the effects of lacunae and canaliculi density, which may also change with aging [56, 57], are not considered. The cement line surrounding osteons has also been shown to play a role in the progression of fractures around or through osteons [48, 58–61]. Its effect on crack propagation seems to partly depend on its composition [23, 48, 58], which is not clear; some studies report high mineralization [62, 63] and others report minimal mineralization [64, 65]. Furthermore, cement lines occupy a very small volume fraction of the bone. Therefore, they were omitted from the model.

Sensitivity values (Figure 7) are the result of output derivatives (Eq. 17) and the expected standard deviation of the inputs: the model’s high sensitivity to post-yield parameters (*β*) arises from the high uncertainty in the values of those parameters, as measured in the coefficients of variation (Table 5). While all mechanical properties were most sensitive to variations in *β* (of either osteon or interstitial fibers), a 10% change in either of these inputs results in only a 1% change in ultimate stress and less than 10% change in toughness. On the other hand, a 10% change in other inputs, such as *α* and osteon volume fraction results in a change of about 20% in both ultimate stress and toughness; since our model predicted a difference of 5% in ultimate strength between the mature and elderly bone cases, these values of 10% and 20% are rather meaningful even if the computed sensitivity is low relative to *α* or β. It is clear that the model’s sensitivity to these *α* and β arises from the high uncertainty in the values of these parameters, due to limited experimental data on post-yield properties of individual microconstituents. For example, one nanoindentation study quantified the aging-related change in the toughness of osteon lamellae [7] but did not report yield stress or plastic modulus. Micropillar experiments produced stress-strain curves of lamellae, but were conducted on dry ovine [66, 67] or bovine specimens [68, 69], which were likely less ductile than fully hydrated tissue [38]. A small amount of data from micropillar tests on human osteons are available, but only for compression rather than tensile loading [19]. Other computational studies, which investigate the crack propagation behavior of cortical bone, have been similarly challenged in defining material properties [13]. For instance, one group estimated the fracture toughness of osteon and interstitial bone tissue based on the difference in their moduli and the homogenized (macroscopic) fracture toughness [23, 70]. The sensitivity of the present model to these parameters suggests that characterizing the post-yield behavior of the microconstituents of Haversian bone is necessary to understand the microstructural basis of the aging-related increase in cortical bone fragility.

The discrepancy between the brittle and elastic-plastic models’ predictions of toughness reveals that decreased toughness is observed only when plasticity is incorporated into fiber material properties. We see in the brittle model that making the interstitial fibers stronger also increases their toughness without also increasing their modulus. Since there is no reported aging-related change in interstitial modulus [12], the increase in strength of the interstitial fibers then increases toughness of the whole bundle. While it is apparent that plasticity within microconstituents plays a key role in the post-yield behavior of cortical bone, a mechanistic understanding of decreased toughness with aging is still lacking.

We find here that aging-related changes other than porosity —in the material properties and relative volume fraction of osteon and interstitial tissue— point toward a microconstitutive basis for increased bone fragility in the elderly population. Our results show that material aging (namely, reduced plasticity) alone and in combination with either porosity increase or morphological aging reduced toughness to a value on par with that seen for the elderly bone. Although porosity has been found to be negatively correlated with yield stress and ultimate stress [5, 6] (and as seen in Case 4, Figure 5), there is agreement that porosity alone cannot fully account for reduction in toughness [6] or fracture toughness [20]. Factors that have been proposed include aging-related changes in microstructure [6, 28] and decrease in plasticity of the bone tissue matrix [5, 28, 31, 71] leading to an accumulation of damage in the form of microcracks [6, 28, 72]. Evidence also points out an aging-associated relationship between decreased toughness and degradation in the integrity [28, 33] as well as dehydration [43, 73] of the mineralized collagen that makes up bone lamellae, which are not explicitly considered in this model. Furthermore, our elastic-plastic model predicted increased toughness when the volume fraction of osteons increased, which reinforces the idea that morphological changes also play a key role in the fragility of cortical bone [6, 74]. This idea is consistent with prior findings that micromechanical properties (i.e., osteon and interstitial hardness and elastic modulus) are sufficient to predict macroscopic stiffness, but not toughness [28]. Furthermore, a study examining the morphology of osteons across young and elderly groups predicted weaker osteons in the elderly group, due to aging-related changes in geometric indices of the osteons (e.g., the percentage of osteon refilling, given by the ratio of total osteon area that is made up by solid bone tissue [25]). When implemented in a numerical study examining three-dimensional micro-morphological changes, changes to the micro-morphology (representative of the osteons bone refilling rate), along with increased porosity, degraded the bulk mechanical properties in the model.

The prediction in Figure 4 showing that the interstitial fibers all rupture before any osteons do is extreme compared to reports in published fractography studies. Those studies show the fracture surfaces to be rough, with osteons protruding up from the fracture plane in ways that suggest comparatively large amounts of plastic strain relative to the interstitial tissue strain. Some histological studies [75–77] of cortical bone that was loaded to failure have noted that most cracks occur in the interstitial tissue, although cracks visibly cross through osteons in some cases [78]. Several studies have pointed out that the relatively brittle interstitial tissue serves as a starting point for initiation and propagation of cracks [79–81]. Other studies report fracture surfaces that demonstrate intact osteons surrounded by interstitial tissue that has completely failed [74, 82–86] (osteon “pullout”), observing plasticity in the osteons prior to rupture. One such study employed scanning electron microscope (SEM) imaging to quantify osteon “pullout” percentage, which measured the ratio of osteon pullout area (the area of debonded osteons that displayed a “pullout” presentation or cavities from which osteons were “pulled out,” characteristic of interstitial failure) to fracture surface area, and showed a decrease in this quantity with aging [84]. While these results seem to be broadly consistent with predictions here that interstitial tissue fails before osteons, they may also be indicative of cement line failure, which is not accounted for by our model. Furthermore, whether cracks propagate through osteons has been shown to depend, in part, on the material properties and geometry of the interface, even if its mechanistic role is not fully understood [23, 48]. While these geometric factors are not considered in the present model, we do predict that osteons, which are considered tougher than interstitial tissue, resist fracture over longer strains than interstitial tissue.

Our results are consistent with the growing understanding that aging-related changes in the morphology and material properties, in addition to porosity, play important roles in aging-induced decreases in strength and toughness of cortical bone. By isolating the mechanical behavior of osteon vs. interstitial tissue, we found that aging-related changes in the plastic behavior of interstitial tissue, as shown in Fig. 4, directly affects the resulting resistance to fracture at the macroscale. The toughness and ultimate strain, in particular, were shown to depend the most on the post-yield properties of interstitial and osteonal bone tissue. Uncertainty due to the lack of data describing the post-yield behavior of interstitial bone tissue contributes substantially to the uncertainty in macroscale predictions of toughness and ultimate strain. It is therefore clear that measurement of these intrinsic material properties will lead to greater insights into the exact nature of aging-related microstructural changes that give rise to aging-related macroscale changes in cortical bone mechanical properties. As cortical bone is a major contributor to whole bone stiffness and strength, further insight into how features besides porosity influence bone fragility will inform our understanding of aging-related fracture.

## A Appendix

### A.1 Type of property distribution

**Figure A.1:**
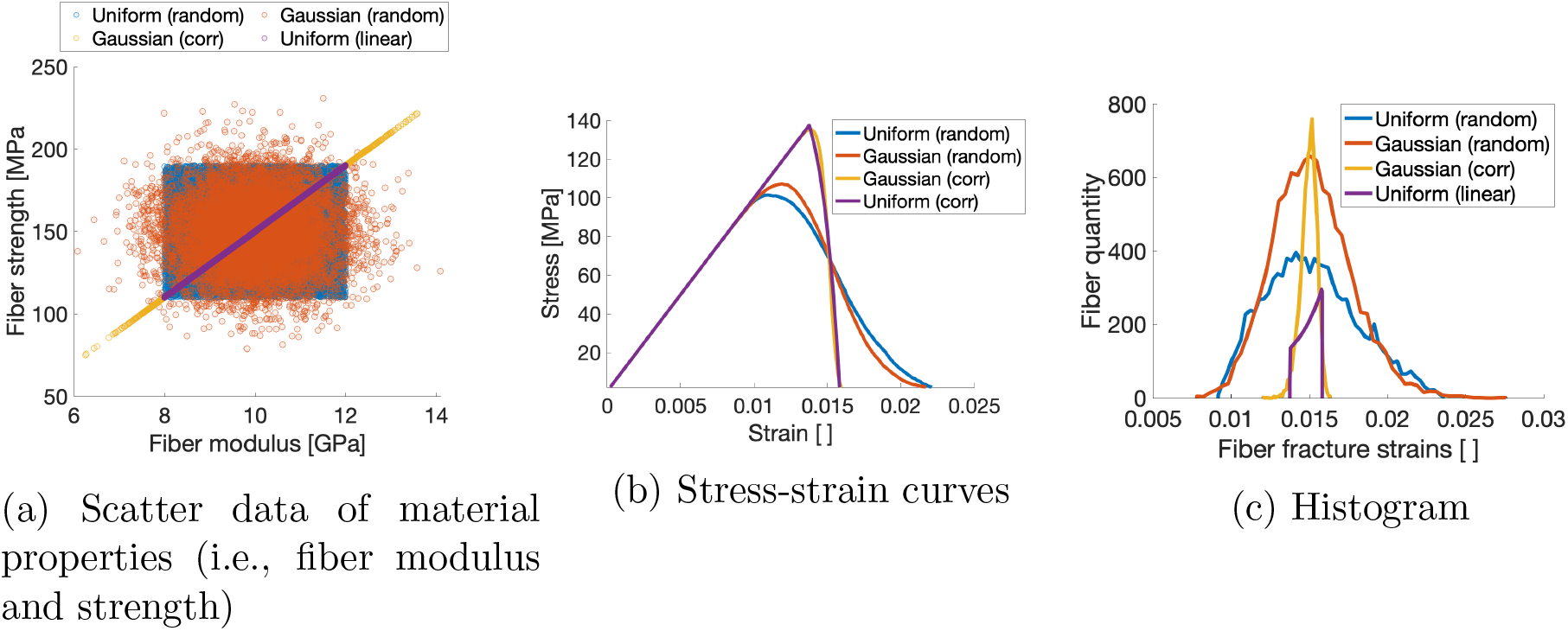
Plots pertaining to 2 distribution types (i.e., uniform and normalized), correlated and randomly sampled.

Accumulation of damage in the bundle is driven by the progressive rupture of individual fibers, which in turn is characterized by the distribution of material properties assigned to the fibers. Thus, the particular type of distribution, and whether the properties are assigned in a manner that makes them correlated with one another (e.g., modulus vs. strength), affects the bundle’s overall mechanical behavior. Prior models feature material properties which are either linearly related [34] or randomly sampled from a uniform distribution [35, 87]. A normal distribution may more accurately reflect the spread in material properties observed in cortical bone. More importantly, the correlation of fiber material properties such as modulus and strength, as shown in (Figure A.1a), also affects model behavior. To investigate this effect, a multivariate Gaussian distribution was constructed with a correlation coefficient (*ρ*) of *1* to correlate the modulus and strength distributions. Simulations of fiber bundles constructed from both correlated and uncorrelated distributions were then conducted to evaluate these potential effects.

For the case of a “correlated” uniform distribution, modulus and strength distributions are defined for monotonically increasing functions of *x* [34].

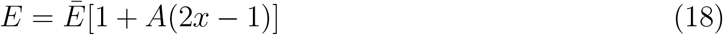

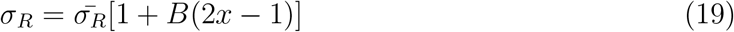

for *x* ∈ [0, 1] (representative of the increasingly strong fibers) where *A* and *B* are characteristic coefficients that each define the extreme values for modulus (E) and rupture strength (*σ_R_*) centered about 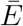 and 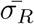, the mean modulus and rupture strength values, respectively.

For a Gaussian distribution in which modulus and rupture strength values are correlated, a multivariate distribution is constructed by first defining a correlation coefficient (*ρ*), which describes the relationship between the two variables. Then, the standard deviation (*σ_x_,σ_y_*) corresponding to each distribution (x,y) is used to construct a covariance matrix (Σ). A built-in function (“mvnrnd”) within MATLAB [88] is implemented to sample an arbitrary number of samples from both distributions according to the specified covariance matrix and distribution means.

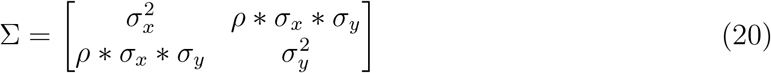

**Table A.1:** Material property distribution parameters - average & standard deviation (Gaussian) and value range (uniform), which define an osteon fiber bundle

|  | Gaussian | Uniform |
| --- | --- | --- |
| Modulus [GPa] | 10 (1) | 8-12 |
| Strength [MPa] | 150 (20) | 110-190 |

The failure behavior of the fiber bundle depended slightly on the shape of the distribution used to define the material properties of the fibers that comprise the bundle, but strongly on their correlation. The ultimate stress of the bundle (Figure A.1b) is increased when the strength of each fiber is correlated with its modulus, regardless of distribution type. Coupled with the decreased ultimate strain and subsequent failure strain (when the bundle stress drops to zero), this results in embrittled behavior. This effect is highlighted when the fracture strains for all fibers in the bundle is considered (Figure A.1c), in which the fibers from the bundle models for which inputted material properties were correlated are narrower than the models in which properties are drawn independently. Though not shown here, we found also that selecting a normal distribution rather than uniform reduces model artifacts when multiple fiber families are incorporated into the model.

### A.2 Convergence analyses: dependence on algorithmic parameters

The number of fibers (“bundle size”) affects how closely the mean and standard deviation of sampled fiber material properties are to the prescribed mean and standard deviation of the material property distribution. To evaluate the effect of the bundle size, simulations with bundles consisting of varying fiber number were conducted; 100 simulations per particular bundle size. Then, the yield stress and ultimate stress for each simulation were computed, as well as their mean and standard deviation for each bundle size. These simulations show that a bundle size on the order of 10^4^ result in a standard deviation of the outputs that is less than 1% of the mean value (Figure A.2).

The strain step size described in Eq. 6 dictates the “time” resolution of the simulation and thus defines how frequently the stress is computed within a given strain range. Since damage in the fiber bundle model is driven by progressive rupture of fibers, the strain resolution may affect the accuracy of the output data; too coarse of a step size could result in a fiber rupturing at a strain larger than would correspond to its material properties. To evaluate the effect of the step size, the yield and ultimate stress were computed for simulations conducted with varying strain step sizes. The model appears to converge at a step size of 10^-4^ (Figure A.3), which indicates that the step size of 10^-5^ used in this study was appropriate.

**Figure A.2:**
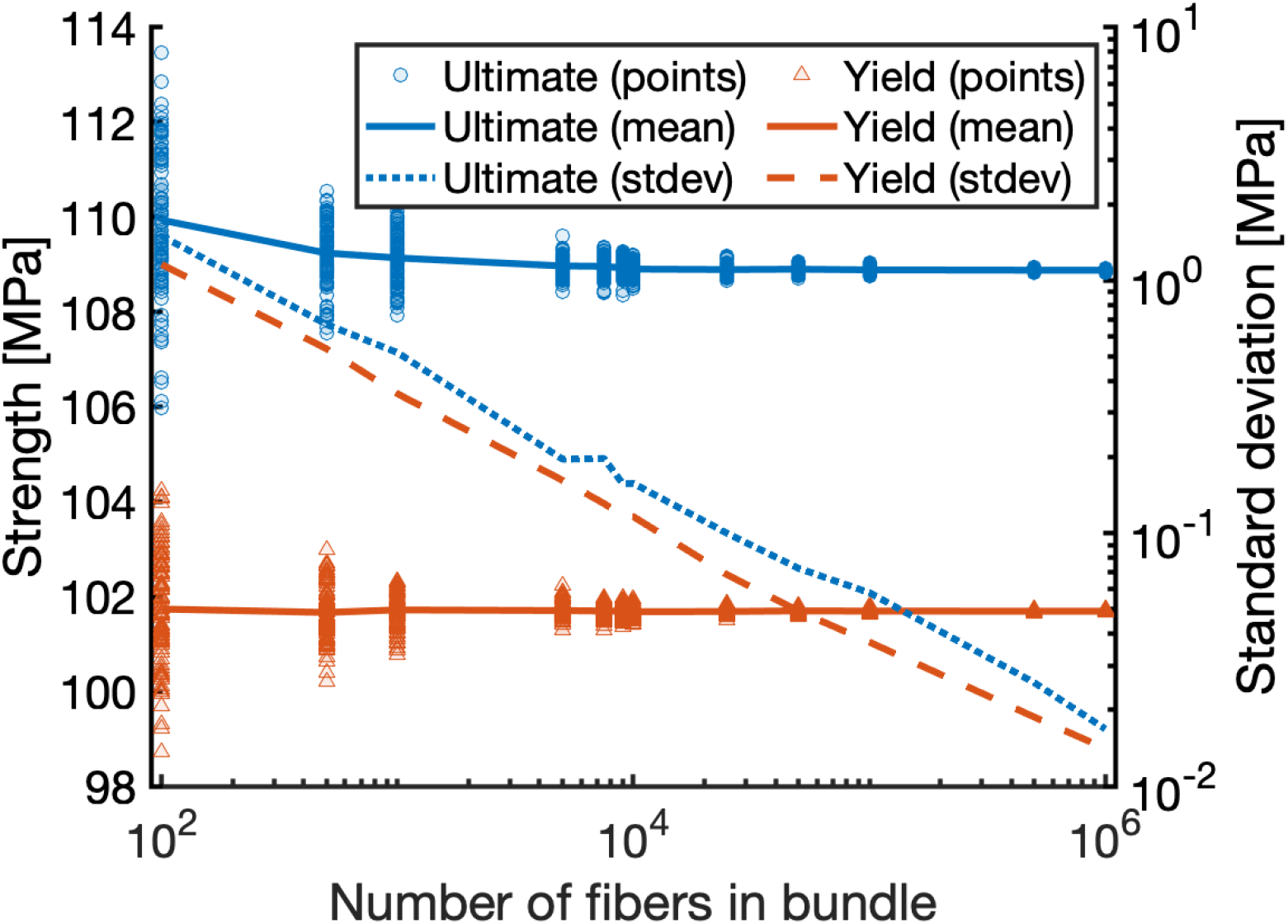
Predicted strength and its standard deviation vs. number of fibers in the bundle. Convergence of ultimate stress (upper line, blue) and yield stress (lower line, red) resulting from randomly sampling model inputs for 100 simulations for each trial number of fibers.

**Figure A.3:**
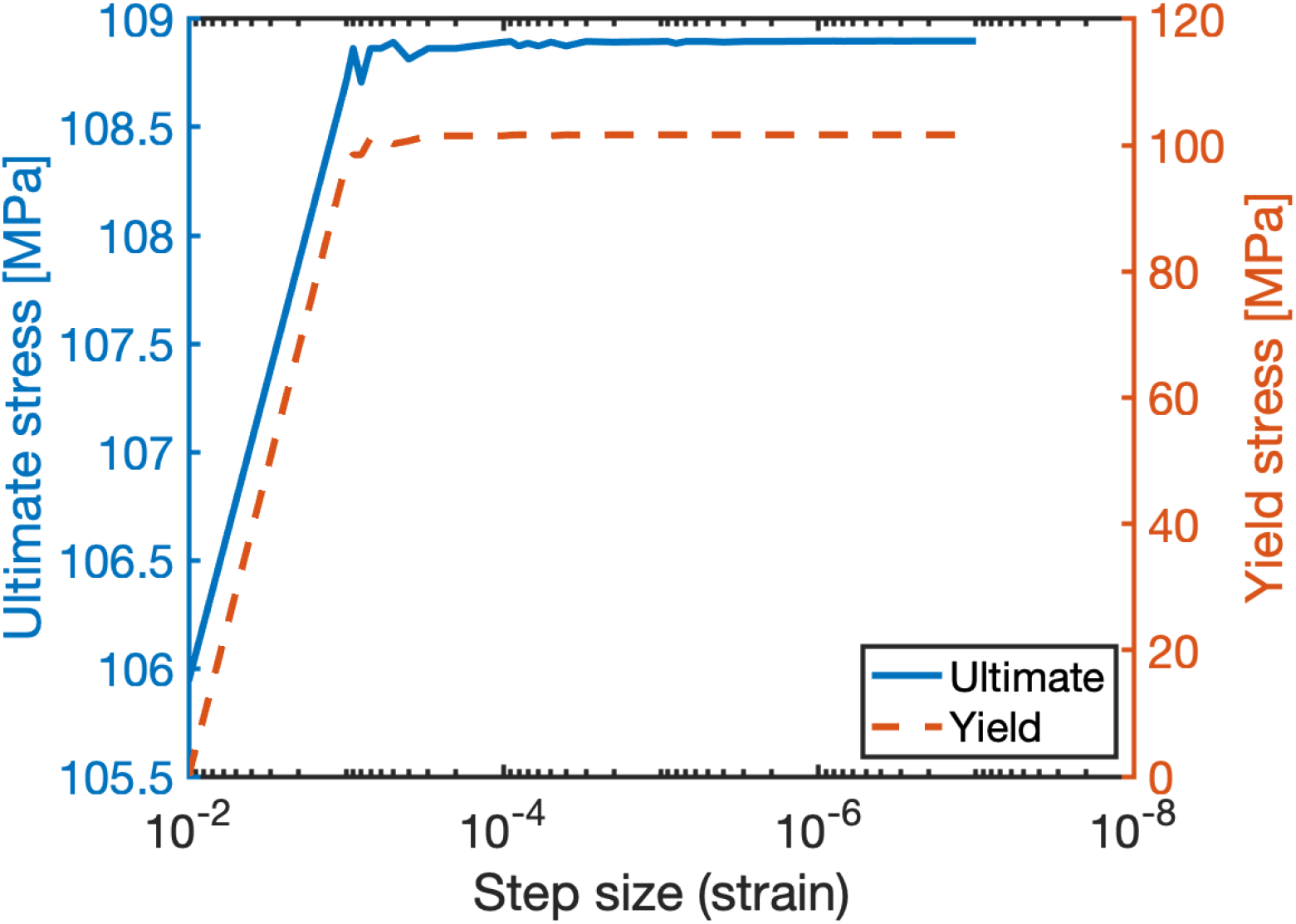
Convergence of ultimate stress (solid) and yield stress (dashed) model outputs from decreasing strain step size.

## Bibliography

[1] Gerold Holzer et al. “Hip fractures and the contribution of cortical versus trabecular bone to femoral neck strength.” In: Journal of bone and mineral research 24.3 (2009), pp. 468–474.

[2] Senthil K Eswaran et al. “Cortical and trabecular load sharing in the human vertebral body.” In: Journal of Bone and Mineral Research 21.2 (2006), pp. 307–314.

[3] Uwe Wolfram and Jakob Schwiedrzik. “Post-yield and failure properties of cortical bone.” In: BoneKEy reports 5 (2016).

[4] Siu L Hui, Charles W Slemenda, C Conrad Johnston, et al. “Age and bone mass as predictors of fracture in a prospective study.” In: The Journal of clinical investigation 81.6 (1988), pp. 1804–1809.

[5] Richard W McCalden, Joseph A McGeough, Michael B Barker, et al. “Age-related changes in the tensile properties of cortical bone. The relative importance of changes in porosity, mineralization, and microstructure.” In: The Journal of bone and joint surgery. American volume 75.8 (1993), pp. 1193–1205.

[6] Mohammad J Mirzaali et al. “Mechanical properties of cortical bone and their relationships with age, gender, composition and microindentation properties in the elderly.” In: Bone 93 (2016), pp. 196–211.

[7] X Wang, YJ Yoon, and H Ji. “A novel scratching approach for measuring age-related changes in the in situ toughness of bone.” In: Journal of biomechanics 40.6 (2007), pp. 1401–1404.

[8] Antonio Ascenzi and Ermanno Bonucci. “The tensile properties of single osteons.” In: The Anatomical Record 158.4 (1967), pp. 375–386.

[9] Yan Wu et al. “Cortical bone mineralization differences between hip-fractured females and controls. A microradiographic study.” In: Bone 45.2 (2009), pp. 207–212.

[10] Z Fan et al. “Anisotropic properties of human tibial cortical bone as measured by nanoindentation.” In: Journal of orthopaedic research 20.4 (2002), pp. 806–810.

[11] Elisa Budyn, Thierry Hoc, and Julien Jonvaux. “Fracture strength assessment and aging signs detection in human cortical bone using an X-FEM multiple scale approach.” In: Computational Mechanics 42.4 (2008), pp. 579–591.

[12] Ji Young Rho et al. “Microstructural elasticity and regional heterogeneity in human femoral bone of various ages examined by nano-indentation.” In: Journal of biomechanics 35.2 (2002), pp. 189–198.

[13] Adel A Abdel-Wahab, Angelo R Maligno, and Vadim V Silberschmidt. “Micro-scale modelling of bovine cortical bone fracture: Analysis of crack propagation and microstructure using X-FEM.” In: Computational Materials Science 52.1 (2012), pp. 128–135.

[14] Jae-Young Rho, Ting Y Tsui, and George M Pharr. “Elastic properties of human cortical and trabecular lamellar bone measured by nanoindentation.” In: Biomaterials 18.20 (1997), pp. 1325–1330.

[15] LP Mullins, MS Bruzzi, and PE McHugh. “Calibration of a constitutive model for the post-yield behaviour of cortical bone.” In: Journal of the Mechanical Behavior of Biomedical Materials 2.5 (2009), pp. 460–470.

[16] CE Hoffler et al. “Heterogeneity of bone lamellar-level elastic moduli.” In: Bone 26.6 (2000), pp. 603–609.

[17] C Edward Hoffler et al. “An application of nanoindentation technique to measure bone tissue lamellae properties.” In: J. Biomech. Eng. 127.7 (2005), pp. 1046–1053.

[18] LP Mullins et al. “Differences in the crack resistance of interstitial, osteonal and trabecular bone tissue.” In: Annals of biomedical engineering 37.12 (2009), pp. 2574–2582.

[19] Caitlyn J Collins et al. “The impact of age, mineralization, and collagen orientation on the mechanics of individual osteons from human femurs.” In: Materialia 9 (2020), p. 100573.

[20] X Wang et al. “Agerelated changes in the mechanical properties of interstitial bone tissues.” In: Proc. 50th Annual Meeting of the Orthopaedic Research Society. 2004.

[21] F Gaynor Evans. “Mechanical properties and histology of cortical bone from younger and older men.” In: The Anatomical Record 185.1 (1976), pp. 1–11.

[22] MKH Malo et al. “Longitudinal elastic properties and porosity of cortical bone tissue vary with age in human proximal femur.” In: Bone 53.2 (2013), pp. 451–458.

[23] Ebrahim Maghami et al. “Fracture behavior of human cortical bone: Role of advanced glycation end-products and microstructural features.” In: Journal of Biomechanics 125 (2021), p. 110600.

[24] Karen M Cooke, Patrick Mahoney, and Justyna J Miszkiewicz. “Secondary osteon variants and remodeling in human bone.” In: The Anatomical Record (2021).

[25] Andreas Bernhard et al. “Micro-morphological properties of osteons reveal changes in cortical bone stability during aging, osteoporosis, and bisphosphonate treatment in women.” In: Osteoporosis International 24.10 (2013), pp. 2671–2680.

[26] P Zioupos and JD Currey. “Changes in the stiffness, strength, and toughness of human cortical bone with age.” In: Bone 22.1 (1998), pp. 57–66.

[27] John D Currey, Kevin Brear, and Peter Zioupos. “The effects of ageing and changes in mineral content in degrading the toughness of human femora.” In: Journal of biomechanics 29.2 (1996), pp. 257–260.

[28] P Zioupos. “Ageing human bone: factors affecting its biomechanical properties and the role of collagen.” In: Journal of Biomaterials Applications 15.3 (2001), pp. 187–229.

[29] Edward D Simmons Jr, KPH Pritzker, and Marc D Grynpas. “Age-related changes in the human femoral cortex.” In: Journal of Orthopaedic Research 9.2 (1991), pp. 155–167.

[30] David ML Cooper et al. “Age-dependent change in the 3D structure of cortical porosity at the human femoral midshaft.” In: Bone 40.4 (2007), pp. 957–965.

[31] JB Phelps et al. “Microstructural heterogeneity and the fracture toughness of bone.” In: Journal of biomedical materials research 51.4 (2000), pp. 735–741.

[32] Xiaodu D Wang et al. “Changes in the fracture toughness of bone may not be reflected in its mineral density, porosity, and tensile properties.” In: Bone 23.1 (1998), pp. 67–72.

[33] X Wang et al. “Age-related changes in the collagen network and toughness of bone.” In: Bone 31.1 (2002), pp. 1–7.

[34] Jean Lemaitre. A course on damage mechanics. Springer Science & Business Media, 2012.

[35] Dusan Krajcinovic. “Damage mechanics.” In: Mechanics of materials 8.2-3 (1989), pp. 117–197.

[36] Hiroshi Yamada, F Gaynor Evans, et al. “Strength of biological materials.” In: (1970).

[37] JC Simo and Thomas J. Hughes. “Motivation. One-Dimensional Plasticity and Viscoplasticity.” In: Computational Inelasticity. New York, NY: Springer New York, 1998, pp. 1–45. ISBN: 978-0-387-22763-4. DOI: 10.1007/0-387-22763-6_1. URL: https://doi.org/10.1007/0-387-22763-6_1.

[38] Anna Faingold et al. “The effect of hydration on mechanical anisotropy, topography and fibril organization of the osteonal lamellae.” In: Journal of Biomechanics 47.2 (2014), pp. 367–372.

[39] JY Rho et al. “Variations in the individual thick lamellar properties within osteons by nanoindentation.” In: Bone 25.3 (1999), pp. 295–300.

[40] Raul Vincentelli and Margarita Grigoroy. “The effect of Haversian remodeling on the tensile properties of human cortical bone.” In: Journal of biomechanics 18.3 (1985), pp. 201–207.

[41] Ozan Akkus et al. “Aging of microstructural compartments in human compact bone.” In: Journal of Bone and Mineral Research 18.6 (2003), pp. 1012–1019.

[42] Andrea Saltelli et al. Global sensitivity analysis: the primer. John Wiley & Sons, 2008.

[43] Jeffry S Nyman et al. “The influence of water removal on the strength and toughness of cortical bone.” In: Journal of biomechanics 39.5 (2006), pp. 931–938.

[44] Albert H Burstein, Donald T Reilly, and Marc Martens. “Aging of bone tissue: mechanical properties.” In: The Journal of bone and joint surgery. American volume 58.1 (1976), pp. 82–86.

[45] Ravi K Nalla et al. “Effect of aging on the toughness of human cortical bone: evaluation by R-curves.” In: Bone 35.6 (2004), pp. 1240–1246.

[46] Amy C Courtney, Wilson C Hayes, and Lorna J Gibson. “Age-related differences in post-yield damage in human cortical bone. Experiment and model.” In: Journal of biomechanics 29.11 (1996), pp. 1463–1471.

[47] Ani Ural and Susan Mischinski. “Multiscale modeling of bone fracture using cohesive finite elements.” In: Engineering Fracture Mechanics 103 (2013), pp. 141–152.

[48] Susan Mischinski and Ani Ural. “Finite element modeling of microcrack growth in cortical bone.” In: Journal of Applied Mechanics 78.4 (2011), p. 041016.

[49] MT Fondrk, EH Bahniuk, and DT Davy. “A damage model for nonlinear tensile behavior of cortical bone.” In: Journal of Biomechanical Engineering 121.5 (1999), pp. 533–541.

[50] D Garcia et al. “A 1D elastic plastic damage constitutive law for bone tissue.” In: Archive of Applied Mechanics 80.5 (2010), pp. 543–555.

[51] Donald T Reilly and Albert H Burstein. “The elastic and ultimate properties of compact bone tissue.” In: Journal of biomechanics 8.6 (1975), pp. 393–405.

[52] Jeffry S Nyman et al. “Age-related factors affecting the postyield energy dissipation of human cortical bone.” In: Journal of orthopaedic research 25.5 (2007), pp. 646–655.

[53] John D Currey. “Some effects of ageing in human Haversian systems.” In: Journal of anatomy 98.Pt 1 (1964), p. 69.

[54] Hayley M Britz et al. “The relation of femoral osteon geometry to age, sex, height and weight.” In: Bone 45.1 (2009), pp. 77–83.

[55] Cheryl Hennig et al. “Does 3D orientation account for variation in osteon morphology assessed by 2D histology?” In: Journal of anatomy 227.4 (2015), pp. 497–505.

[56] Petar Milovanovic et al. “Osteocytic canalicular networks: morphological implications for altered mechanosensitivity.” In: ACS nano 7.9 (2013), pp. 7542–7551.

[57] Jiarui Cui et al. “Osteocytes in bone aging: Advances, challenges, and future perspectives.” In: Ageing research reviews (2022), p. 101608.

[58] Anna Gustafsson et al. “Crack propagation in cortical bone is affected by the characteristics of the cement line: a parameter study using an XFEM interface damage model.” In: Biomechanics and modeling in mechanobiology 18.4 (2019), pp. 1247–1261.

[59] Eugenio Giner et al. “Calculation of the critical energy release rate Gc of the cement line in cortical bone combining experimental tests and finite element models.” In: Engineering Fracture Mechanics 184 (2017), pp. 168–182.

[60] Sabah Nobakhti, Georfes Limbert, and Philipp J. Thurner. “Cement lines and interlamellar areas in compact bone as strain amplifiers - Contributors to elasticity, fracture toughness and mechanotransduction.” In: Journal of the Mechanical Behavior of Biomedical Materials 29 (2014), pp. 235–251.

[61] X Neil Dong, Xiaohui Zhang, and X Edward Guo. “Interfacial strength of cement lines in human cortical bone.” In: Molecular & Cellular Biomechanics 2.2 (2005), p. 63.

[62] John G. Skedros et al. “Cement lines of secondary osteons in human bone are not mineral-deficient: New data in a historical perspective.” In: The Anatomical Record 286A.1 (2005), pp. 781–803.

[63] Petar Milovanovic et al. “Bone tissue aging affects mineralization of cement lines.” In: Bone 110 (2018), pp. 187–193.

[64] Timothy Montalbano and Gang Feng. “Nanoindentation characterization of the cement lines in ovine and bovine femurs.” In: Journal of Materials Research 26.8 (2011), pp. 1036–1041.

[65] David B Burr, Mitchell B Schaffler, and Richard G Frederickson. “Composition of the cement line and its possible mechanical role as a local interface in human compact bone.” In: Journal of biomechanics 21.11 (1988), pp. 939–945.

[66] Daniele Casari et al. “Microtensile properties and failure mechanisms of cortical bone at the lamellar level.” In: Acta biomaterialia 120 (2021), pp. 135–145.

[67] Jakob Schwiedrzik et al. “In situ micropillar compression reveals superior strength and ductility but an absence of damage in lamellar bone.” In: Nature materials 13.7 (2014), pp. 740–747.

[68] Krzysztof W Luczynski et al. “Extracellular bone matrix exhibits hardening elastoplasticity and more than double cortical strength: Evidence from homogeneous compression of non-tapered single micron-sized pillars welded to a rigid substrate.” In: Journal of the mechanical behavior of biomedical materials 52 (2015), pp. 51–62.

[69] Jose A Robles-Linares et al. “Machining-induced thermal damage in cortical bone: Necrosis and micro-mechanical integrity.” In: Materials & Design 197 (2021), p. 109215.

[70] Timothy O Josephson et al. “Computational study of the mechanical influence of lacunae and perilacunar zones in cortical bone microcracking.” In: Journal of the Mechanical Behavior of Biomedical Materials 126 (2022), p. 105029.

[71] Yohann Bala et al. “Respective roles of organic and mineral components of human cortical bone matrix in micromechanical behavior: an instrumented indentation study.” In: Journal of the mechanical behavior of biomedical materials 4.7 (2011), pp. 1473–1482.

[72] Timothy L Norman et al. “Influence of microdamage on fracture toughness of the human femur and tibia.” In: Bone 23.3 (1998), pp. 303–306.

[73] Mathilde Granke, Mark D Does, and Jeffry S Nyman. “The role of water compartments in the material properties of cortical bone.” In: Calcified tissue international 97.3 (2015), pp. 292–307.

[74] DD Moyle and RW Bowden. “Fracture of human femoral bone.” In: Journal of biomechanics 17.3 (1984), pp. 203–213.

[75] David B Burr. “Stress concentrations and bone microdamage: John Currey’s contributions to understanding the initiation and arrest of cracks in bone.” In: Bone 127 (2019), pp. 517–525.

[76] W Bonfield and JC Behiri. “Fracture toughness of natural composites with reference to cortical bone.” In: Composite Materials Series. Vol. 6. Elsevier, 1989, pp. 615–635.

[77] John G Skedros, Mark W Mason, and Roy D Bloebaum. “Differences in osteonal micromorphology between tensile and compressive cortices of a bending skeletal system: indications of potential strain-specific differences in bone microstructure.” In: The Anatomical Record 239.4 (1994), pp. 405–413.

[78] Susan Mischinski and Ani Ural. “Interaction of microstructure and microcrack growth in cortical bone: a finite element study.” In: Computer methods in biomechanics and biomedical engineering 16.1 (2013), pp. 81–94.

[79] Todd M Boyce et al. “Damage type and strain mode associations in human compact bone bending fatigue.” In: Journal of Orthopaedic Research 16.3 (1998), pp. 322–329.

[80] MB Schaffler, K Choi, and C Milgrom. “Aging and matrix microdamage accumulation in human compact bone.” In: Bone 17.6 (1995), pp. 521–525.

[81] Timothy L Norman and Z Wang. “Microdamage of human cortical bone: incidence and morphology in long bones.” In: Bone 20.4 (1997), pp. 375–379.

[82] Francis W Cooke, Howard Zeidman, and Stephen J Scheifele. “The fracture mechanics of bone—another look at composite modeling.” In: Journal of Biomedical Materials Research 7.3 (1973), pp. 383–399.

[83] G Corondan and WL Haworth. “A fractographic study of human long bone.” In: Journal of biomechanics 19.3 (1986), pp. 207–218.

[84] LP Hiller et al. “Osteon pullout in the equine third metacarpal bone: effects of ex vivo fatigue.” In: Journal of orthopaedic research 21.3 (2003), pp. 481–488.

[85] DD Moyle, JW Welborn III, and FW Cooke. “Work to fracture of canine femoral bone.” In: Journal of Biomechanics 11.10-12 (1978), pp. 435–440.

[86] MH Pope and JO Outwater. “The fracture characteristics of bone substance.” In: Journal of Biomechanics 5.5 (1972), pp. 457–465.

[87] Dusan Krajcinovic, Jordan Trafimow, and Dragoslav Sumarac. “Simple constitutive model for a cortical bone.” In: Journal of biomechanics 20.8 (1987), pp. 779–784.

[88] MATLAB. version 7.10.0 (R2010a). Natick, Massachusetts: The MathWorks Inc., 2022.

